# Short stop is a gatekeeper at the ring canals of *Drosophila* ovary

**DOI:** 10.1101/2020.12.09.418046

**Authors:** Wen Lu, Margot Lakonishok, Vladimir I. Gelfand

## Abstract

Microtubules and actin filaments are two major cytoskeletal components essential for a variety of cellular functions. Spectraplakins are a family of large cytoskeletal proteins cross-linking microtubules and actin filaments among other components. In this study, we aim to understand how Short stop (Shot), the single *Drosophila* spectraplakin, coordinates microtubules and actin filaments for oocyte growth. The oocyte growth completely relies on the acquisition of cytoplasmic materials from the interconnected sister cells (nurse cells), through ring canals, cytoplasmic bridges that remained open after incomplete germ cell division. Given the open nature of the ring canals, it is unclear how the direction of transport through the ring canal is controlled. Here we show that Shot controls the directionality of flow of material from the nurse cells towards the oocyte. Knockdown of *shot* changes the direction of transport of many types of cargo through the ring canals from unidirectional (toward the oocyte) to bidirectional, resulting in small oocytes that fail to grow over time. In agreement with this flow-directing function of Shot, we find that it is localized at the asymmetric actin fibers adjacent to the ring canals at the nurse cell side, and controls the uniform polarity of microtubules located in the ring canals connecting the nurse cells and the oocyte. Together, we propose that Shot functions as a gatekeeper directing the material flow from the nurse cells to the oocyte, via organization of microtubule tracks.

## INTRODUCTION

Microtubules and actin filaments are two fundamental cytoskeletal components of all eukaryotic cells. They are essential for multiple key functions of a cell, such as cell division, cell migration, cargo transport, morphogenesis and compartmentation/polarization. Coordination of microtubules and actin filaments is vital for these various cellular functions. Yet full understanding of microtubule-actin crosstalk is still lacking.

Spectraplakins are a family of large cytoskeletal linker proteins that are evolutionarily conserved across the animal kingdom. Spectraplakins are unique in their ability to associate with all three cytoskeletal networks: F-actin, microtubules and intermediate filaments. They all contain N-terminal calponin homology (CH) domains for actin binding (ABD), C-terminal EF motif, GAS2 domain and C-terminal tail containing plus-end tracking SxIP motifs (EGC) for microtubule interaction, bridged by a plakin domain and a long rod-like domain composed of spectrin repeats [1, 5]. Short stop (Shot) is the single *Drosophila* spectraplakin, coordinating and moderating the interactions between F-actin and microtubules via the N-terminal ABD domain and C-terminal EGC domain, respectively (Figure 1A) [6, 7]. *Drosophila* Shot has been shown to be involved in the regulation of cytoskeletal network interaction in many cell types [8]. For instance, Shot controls microtubule organization and regulates filopodia formation in neurites and is thus essential for axon extension [6, 7, 9-11]. Furthermore, Shot plays a critical role in multiple cell shape changes and developmental morphogenesis events, such as tracheal tube fusion [12, 14], epithelia cell-cell adhesion [15], foregut development [16], photoreceptor morphogenesis [17], salivary gland tube formation [17], muscle myonuclear shape maintenance [18] and dorsal closure [19].

**Figure 1.**
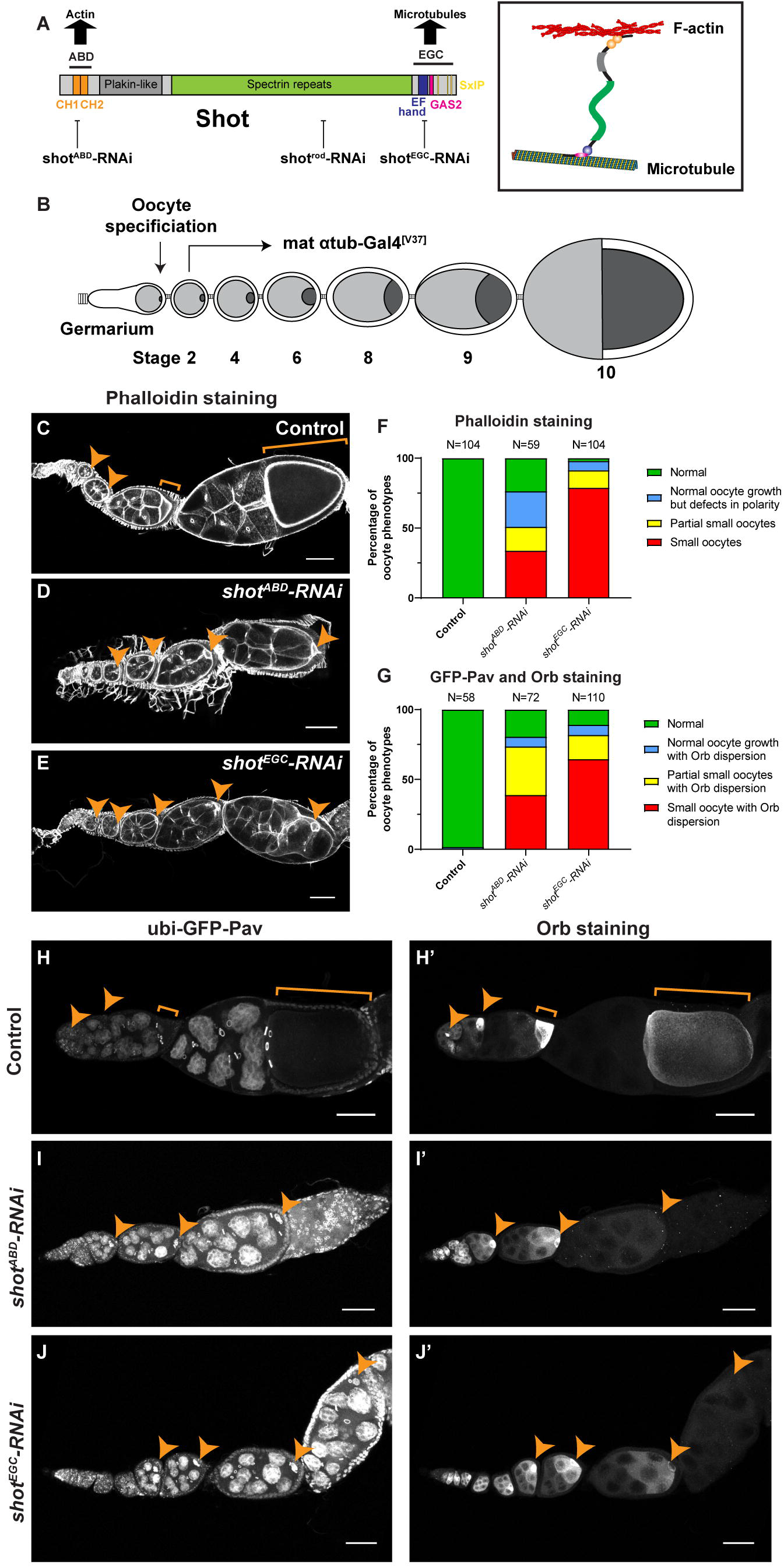
Shot is required for Drosophila oocyte growth. (A) A diagram of Shot multiple domains and Shot crosslinking activity of microtubules and F-actin. Three independent *shot-RNAi* lines were used in this study, targeting ABD, Rod and EGC domains, respectively. (B) A schematic illustration of *Drosophila* oogenesis in one ovariole and the *shot*-RNAi knockdown strategy to bypass the requirement of Shot in oocyte specification. Oocyte is shown in darker grey, while nurse cells are represented in lighter grey in the egg chambers. The *mat* α*tub-Gal4^[V37]^* line starts the expression in stage 2-3 egg chambers, after the completion of oocyte specification. (C-E) Representative images of Rhodamine-conjugated phalloidin staining in control (*mat* α*tub-Gal4^[V37]^/+*), *shot^ABD^-RNAi* (*mat* α*tub-Gal4^[V37]^/UAS-shot^ABD^-RNAi*), and *shot^EGC^-RNAi* (*mat* α*tub-Gal4^[V37]^/UAS-shot^EGC^-RNAi*). See also in Video 1. (F) Summary of phalloidin staining phenotypes in control, *shot^ABD^-RNAi* and *shot^EGC^-RNAi*. (G) Summary of GFP-Pav labeling and Orb staining phenotypes in control, *shot^ABD^-RNAi* and *shot^EGC^-RNAi*. (H-J’) Representative images of GFP-Pav labeling (H-J) and Orb staining (H’-J’) in control (*ubi-GFP-Pav/+; mat* α*tub-Gal4^[V37]^/+*)*, shot^ABD^-RNAi* (*ubi-GFP-Pav/+; mat* α*tub-Gal4^[V37]^/UAS-shot^ABD^-RNAi*), and *shot^EGC^-RNAi* (*ubi-GFP-Pav/+; mat* α*tub-Gal4^[V37]^/UAS-shot^EGC^-RNAi*). See also in Video 2. Oocytes are highlighted by orange arrowheads or brackets. Scale bars, 50 μm.

The *Drosophila* oocyte is the largest cell in a fruit fly. An oocyte is first specified among 16-interconnected cyst cells with a diameter of several micrometers, and grows to a full-size of several hundred micrometers, increasing its size by more than a hundred thousand times to prepare for future embryogenesis [20]. Remarkably, the *Drosophila* oocyte is mostly transcriptionally silent throughout oogenesis [21], and its drastic growth is completely dependent on the acquisition of organelles, mRNA, and proteins from the interconnected nurse cells, through ring canals, the intercellular cytoplasmic channels remained after incomplete cytokinesis [22]. Therefore, it is critical to understand what controls the direction of cytoplasmic transport from the nurse cells to the oocyte to support the oocyte growth. Given that microtubules and actin are both present at the nurse cell-oocyte ring canals [23, 25], Shot, the microtubule-actin crosslinker, appears to be an interesting candidate that could regulate cytoplasmic transfer to the oocyte.

Previous studies have shown that Shot is essential for *Drosophila* oogenesis. At early stages, Shot is required for oocyte specification in 16-cell cysts via association of microtubules with fusome [26], a membranous structure in interconnected germline cysts that contains several actin-related cytoskeletal proteins, such as adducin-like Hts and α-spectrin [27]. In mid-oogenesis, Shot links minus-ends of microtubules to the anterior and lateral actin cortex via a minus-end binding protein Patronin, and therefore is essential for the anterior-posterior microtubule gradient formation within the oocyte [28]. This Shot-dependent microtubule gradient is required to control oocyte nucleus translocation and axis determination for future embryos [29, 30]. However, little is known about whether Shot plays a role in oocyte growth because of the fact that *shot* null mutant fails to specify the oocyte [26].

In this study, we use a germline specific Gal4 that drives *shot-RNA*i after oocyte specification and show that Shot is essential for the oocyte growth. Live cell microscopy demonstrates that Shot controls directionality of transport of multiple cargoes through the nurse cell-oocyte ring canals. In the wild-type egg chambers this transport is unidirectional, but after Shot knockdown the transport becomes bidirectional and thus oocyte growth is stalled. Consistent with the fact that Shot controls transport directionality, we discover that Shot is asymmetrically localized at the ring canals connecting nurse cells with the oocyte. It is found on actin fibers that form baskets on the nurse cell side of the ring canals. Furthermore, Shot controls the orientation of microtubules present inside the ring canals: while in the wild-type microtubules are orientated predominantly with minus-ends towards the oocyte, knocking down Shot results in a mixed polarity of ring canal microtubules. We propose that Shot organizes microtubules in the ring canals, allowing the minus-end-directed motor, cytoplasmic dynein, to transport multiple cytoplasmic components from the nurse cells to the oocyte, which is required for the oocyte rapid growth.

## RESULTS

### Shot is essential for oocyte growth

Shot, a single spectraplakin in *Drosophila*, is essential for oogenesis. Each *Drosophila* ovary is composed of 15~20 individual developmental “assembly lines”, called ovarioles. Oogenesis starts in the most anterior structure of each ovariole, the germarium, and one oocyte is specified within a cyst of 16-interconected germline cells. The oocyte, together with 15-interconnected germline cells, nurse cells, gets encapsulated by a mono-layer of somatic follicle cells and become an egg chamber [20](Figure 1B). Germline clone mutant for *shot*^*3*^ [6], a protein-null allele of *shot*, leads to failure of oocyte specification, shown by lack of an concentrated oocyte marker, Orb (oo18 RNA-binding protein) [31] (Supplementary Figure 1A-1B), consistent with a previous report [26].

In order to avoid the early oocyte specification defects, we used a maternal α-tubulin-Gal4 (*mat* α*tub-Gal4*^*[V37]*^) to drive *shot-RNAi*. This Gal4 drives expression starting in stage 2-3 egg chambers after completion of cell divisions and oocyte specification [32, 34], thus bypassing the requirement of Shot for oocyte specification (Figure 1B). We use three different RNAi lines targeting the N-terminus (*shot^ABD^-RNAi*), C-terminus (*shot^EGC^-RNAi*), or the middle rod domain (*shot^Rod^-RNAi)* of *shot*, respectively (Figures 1A). The depletion of Shot after oocyte specification by maternal α-tubulin-Gal4 still allows oocyte specification to occur in early egg chambers, but causes striking oocyte growth defects (referred as “small oocyte phenotype”). These oocytes (identified as the single germline cell with four ring canals, and a non-polyploid nucleus, by phalloidin staining or GFP-tagged kinesin-6/Pavarotti labeling[35]) remain small and fail to grow over-time (Figure 1C-1E and 1H-J’; Videos 1 and 2). All three RNAi lines driven by maternal α-tubulin-Gal4 display the small oocyte phenotype with slight differences in penetrance (Figures 1F-1G and 2A-2B; Supplementary Figure 2A-2F’), indicating that this phenotype is specific to *shot* knockdown. Therefore, we conclude that Shot is essential for oocyte growth.

**Figure 2.**
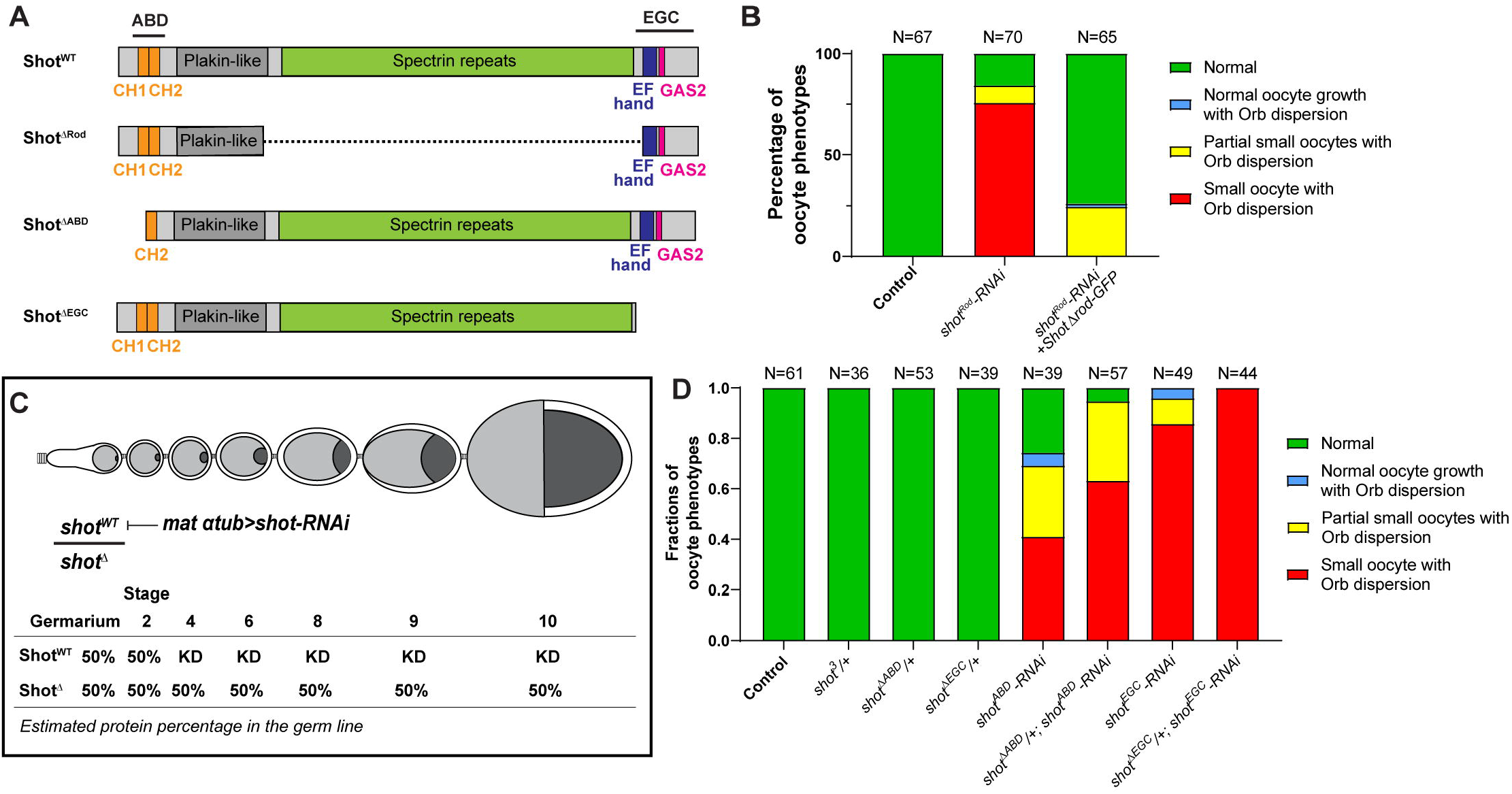
Actin binding and microtubule interacting domains of Shot are essential for oocyte growth. (A) Diagrams of the full-length Shot and truncated mutants. (B) Summary of Orb and phalloidin staining phenotypes in control (*mat* α*tub-Gal4^[V37]^/+*), *shot^Rod^-RNAi* (*mat* α*tub-Gal4^[V37]^/UASp-shot^Rod^-RNAi*), and *shot^Rod^-RNAi* +*shot*Δ*Rod-GFP* (*UASt-shot.L(A)*Δ*Rod-GFP/+; mat* α*tub-Gal4^[V37]/^UASp-shot^Rod^-RNAi*). (C) A schematic illustration of knockdown of wild-type Shot by *shot-RNAi* in *shot* truncated mutant heterozygous background. KD, knockdown. (D) Summary of Orb and phalloidin staining in listed phenotypes. Unlike one copy of *shot^WT^*, one copy of *shot*^Δ*ABD*^ or *shot*^Δ*EGC*^ is unable to drive normal oocyte growth.

### Actin binding domain and microtubule interacting domains of Shot are required for oocyte growth

Shot is a giant cytoskeletal protein, carrying the N-terminal actin binding domain (ABD, composed of CH1 and CH2 domain) and the C-terminal microtubule interacting domain (EGC, composed of EF hand motif, GAS2 domain and C-terminal tail with SxIP motifs) connected by a long rod-like domain composed of spectrin repeats (Figure 1A) [4, 6, 7, 36]. Therefore, we decided to determine which domain is essential for oocyte growth. The long rod domain of spectrin-repeats is essential for intramolecular head-to-tail auto-inhibition of Shot [36]. We first tested whether the rod domain is required to drive the oocyte growth. The *shot-RNAi* targeting the rod region (*shot^Rod^-RNAi*) (Figure 1A) caused majority of the ovarioles displays oocyte growth defects (Figure 2B). The maternal expression of the Shot^ΔRod^ construct lacking the spectrin repeats (Figure 2A) rescued the “small oocyte” phenotype, resulting in most of the ovarioles with normal oocyte growth and concentrated Orb staining (Figure 2B). This indicates that the spectrin repeats are indeed dispensable for normal oocyte growth.

Next, we tested whether ABD and EGC domains are required for oocyte growth. As the Shot mutant lacking the EGC domain (*shot*^Δ*EGC*^) is not homozygous viable [19], we induced germline clones that are homozygous of *shot*^Δ*EGC*^. We found that, similar to *shot^3^* germline clones, *shot*^Δ*EGC*^ mutant clones fail to specify oocytes, shown by the Orb staining (Supplementary Figure 1C-1D). In this case, we cannot determine whether the microtubule interacting domain is essential for oocyte growth due to the complete absence of oocyte specification in the *shot*^Δ*EGC*^ mutant clones. Therefore, we took advantage of the fact that the maternal α-tubulin-Gal4 drives RNAi expression after oocyte specification and combined it with heterozygous truncated mutants that lacks either ABD domain (*shot*^Δ*ABD*^/*shot^WT^*) or EGC domain (*shot*^Δ*EGC*^/*shot^WT^*). We chose the RNAi that only specifically knocks down wild-type *shot*, leaving the truncated *shot* mutant intact (*shot^ABD^-RNAi in shot*^Δ*ABD*^/*shot^WT^* background, and *shot^EGC^-RNAi in shot*^Δ*EGC*^/*shot^WT^* background, respectively) (Figure 2C). In this scenario, a single copy of the wild-type *shot* would specify oocyte fate properly before it gets knocked down by *shot-RNAi* driven by maternal α-tubulin-Gal4 (starting at stage 2-3), which allows us to determine whether the single copy of truncated *shot* mutant could drive oocyte growth after stage 3 (Figure 2C). First of all, we confirmed that oogenesis is completely normal with one single copy of wild-type *shot* (*shot^3^*/*shot^WT^*, *shot*^Δ*ABD*^/*shot*^*WT*^ and *shot*^Δ*EGC*^/*shot^WT^*), excluding the possibility of haploinsufficiency (Figure 2D). Then comparing the *shot-RNAi* in wild-type *shot* background versus in the heterozygous background of *shot* truncated mutant that is insensitive to the *shot-RNAi*, we found that neither one copy of *shot*^Δ*ABD*^ nor one copy of *shot*^Δ*EGC*^ is able to drive oocyte growth (Figure 2D). Together, these data indicated that both the actin binding and the microtubule interacting domains are essential for Shot’s function in promoting oocyte growth, while the central domain is dispensable.

### Shot defines the direction of cargo transport through the nurse cell-oocyte ring canals

The *Drosophila* oocyte, remaining transcriptionally silent throughout most of the oogenesis, completely replies on its sister nurse cells for providing mRNA, proteins and organelles for its growth. The small oocyte phenotype we observed in *shot-RNAi* suggested some defects in cargo transport from the nurse cells to the oocyte. Furthermore, we noticed that in *shot-RNAi* Orb staining is correctly concentrated in the oocyte in early-stage egg chambers; however, the Orb staining becomes more dispersed and eventually lost in the oocytes (Figure 1H’-J’; Video 2). This suggested that the oocyte fails to retain ooplasmic components after receiving them from the sister nurse cells. Therefore, we decided to examine the role of Shot in the transfer of materials from nurse cells to the oocyte. We selected four types of cargoes that are important for oocyte function: Golgi units, ribonucleoprotein particles (RNPs), mitochondria, and lipid droplets (LDs), and studied the role of *shot* in the direction of transport of these components through the ring canals connecting nurse cells and the oocyte.

The Golgi is in the center of secretory pathway and membrane trafficking, which are vital for oocyte development [24, 37]. Using a RFP-tagged Golgi line [38], we were able to visualize robust Golgi unit movements within the nurse cells, and between the nurse cells and the oocyte (Video 3). As previously documented, the vast majority of Golgi units are transported from the nurse cells towards the oocyte through the ring canals (Figure 3A and 3C) [24]. However, in *shot-RNAi* mutant, the directionality of Golgi transport is completely disrupted. We observed frequent reversal of Golgi transport, when the Golgi units move through the ring canals in the opposite direction, from the oocyte back to nurse cells (Figure 3B-3C; Video 3).

**Figure 3.**
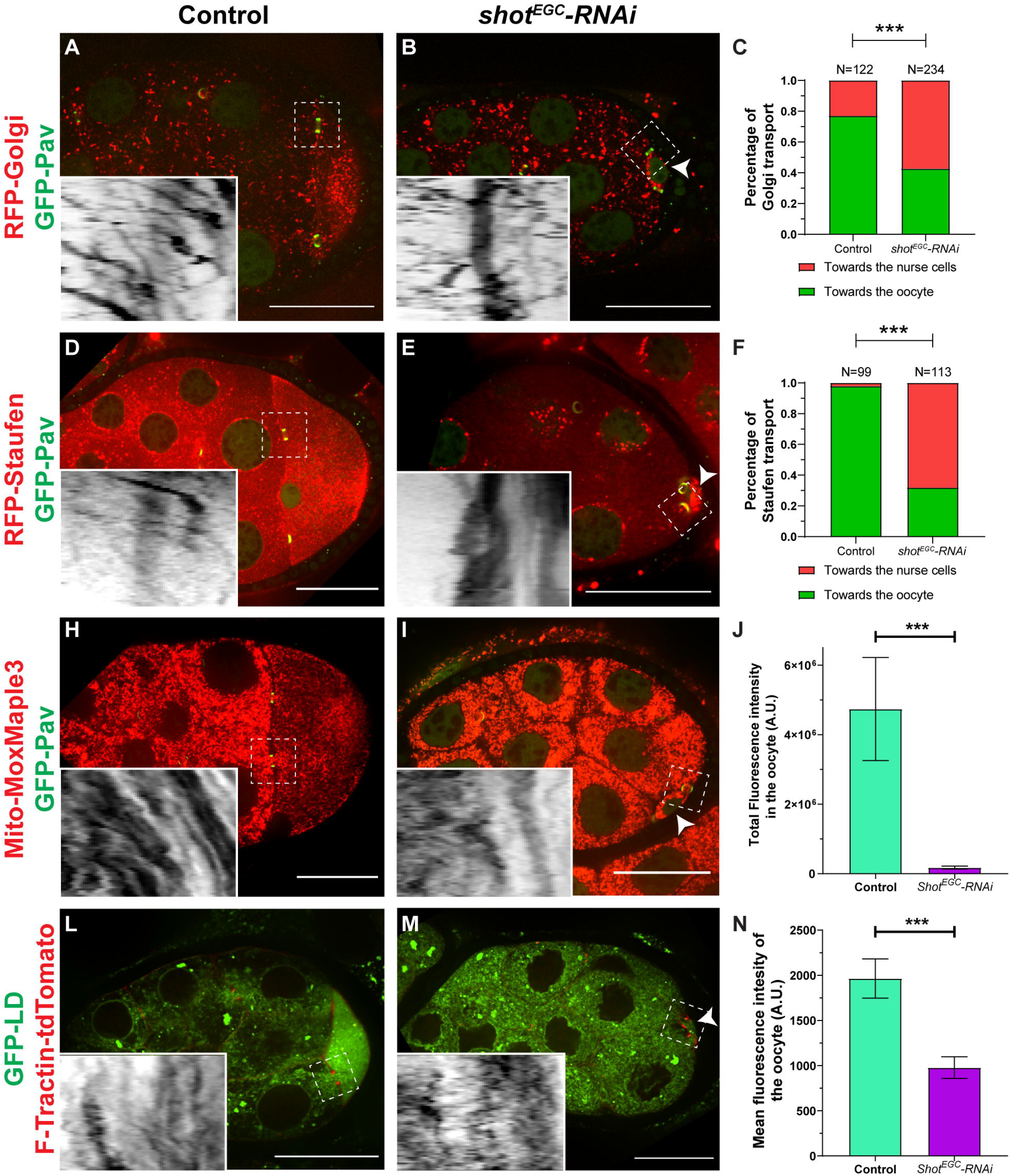
Shot controls directionality of cargo transport from the nurse cells to the oocyte. (A-C) Golgi transport at the nurse cell-oocyte ring canals in control (A) and in *shot-RNAi* (B). Golgi are labeled with RFP-tagged human galactosyltransferase (GalT) (RFP-Golgi). (C) Quantification of Golgi transport directions in control and in *shot-RNAi*. Chi-square test, p-value < 0.00001 (***). See also in Video 3. (D-F) Staufen RNP transport at the nurse cell-oocyte ring canals in control (D) and in *shot-RNAi* (E). Staufen RPNs are labeled with RFP-tagged Staufen (RFP-Staufen). (F) Quantification of Staufen transport directions in control and in *shot-RNAi*. Chi-square test, p-value < 0.00001 (***). See also in Video 4. (H-J) Mitochondria transport at the nurse cell-oocyte ring canals in control (H) and in *shot-RNAi* (I). Mitochondria are labeled with Mito-MoxMaple3 (red channel, after global photoconversion). (J) Quantification of total mitochondria fluorescence intensity (mean ± 95% confidence interval) in control (N=18) and in *shot-RNAi* (N=22) oocytes. Mann-Whitney test, p-value < 0.0001 (***). See also in Video 6. (L-N) Transport of lipid droplets at the nurse cell-oocyte ring canals in control (L) and in *shot-RNAi* (M). Lipid droplets are labeled with GFP-LD. (N) Quantification of average lipid droplet fluorescence intensity (mean ± 95% confidence interval) in control (N=33) and in *shot-RNAi* (N=28) oocytes. Mann-Whitney test, p-value < 0.0001 (***). See also in Video 7. Left side: the nurse cell; right side, the oocyte; small oocytes in *shot-RNAi* are pointed with the white arrowheads; ring canals are labeled with either GFP-Pav (A-B, D-E, H-I) or F-Tractin-tdTomato (L-M); Insets, inverted kymographs were created along a ~3.7μm-width line from the nurse cell to the oocyte through the ring canals in the white dashed box area; scale bars, 50 μm.

During mid-oogenesis, mRNA localization at specific regions of the *Drosophila* oocyte specify the future embryonic axes. Particularly, *osk*/Staufen RNPs are produced in nurse cells, transported into the oocyte and accumulated at its posterior pole specifying posterior determination [39]. Here we examined transport of *osk*/Staufen RNP particles using RFP-Staufen as a marker [30, 40]. In the control egg chambers, transport of RFP-Staufen through the nurse cell-oocyte ring canals is unidirectional (Figure 3D and 3F; Video 4). As in the case of the Golgi units, knockdown of *shot* dramatically changes the transport, resulting in large number of Staufen particles moving from the oocyte back into the nurse cells (Figure 3E-3F; Video 4).

Maternally loaded mitochondria play an essential role in *Drosophila* embryogenesis and germ cell formation. Mitochondria are transported from sister nurse cells and concentrated in the oocyte [41, 42]. To examine mitochondria movement, we employed a newly developed photoconvertible probe (MoxMaple3) [43] targeted to mitochondria (Mito-MoxMaple3, see more details in Materials and Methods). First, we performed local photoconversion of mitochondria either in posterior-most nurse cells or in oocytes and tracked this specific population of mitochondria. We observed red mitochondria photoconverted in the nurse cells moving into the oocyte, but no red mitochondria photoconverted in the oocyte entering the nurse cells (Video 5). Furthermore, after global photoconversion red mitochondria move actively within the nurse cells, and pass through the ring canals from the nurse cell to the oocyte (Figure 3H; Video 6), similar to a previous report using YFP-tagged mitochondria [44]. With *shot-RNAi*, this strict unidirectionality is severely disrupted. We observed very fast but chaotic bidirectional movement through the ring canals (Figure 3I; Video 6), resulting in a reduced number of mitochondria in the oocyte (Figure 3J).

Lipid droplets are generated in the nurse cells and accumulated in the oocyte starting mid-oogenesis. Lipid droplets are believed to be the major energy source for developing embryos as well as an important source for generating membrane components and signaling molecules [45]. Here we used a GFP-tagged lipid droplet domain of *Drosophila* protein Klar (GFP-LD) that is known to target lipid droplets in *Drosophila* germline cells [46]. We found that lipid droplets are transported from the nurse cells to the oocyte through the ring canals in an orderly and consistent fashion in control (Figure 3L; Video 7). In contrast, in *shot-RNAi* egg chambers, lipid droplets move bidirectionally between nurse cells and the oocyte (Figure 3M; Video 7) and fail to concentrate in the oocyte (Figure 3N).

Collectively, these data demonstrate that Shot controls the directionality of material flow from the nurse cells to the oocyte. Lack of Shot results in random transport of cargoes between nurse cells and the oocyte, likely underlying the oocyte growth arrest.

### Localization of Shot on the nurse cell side of the ring canals

Having confirmed that Shot controls the directionality of transport between nurse cells and the oocyte, we decided to examine Shot localization around the ring canal region.

As previously described, actin filaments form asymmetric baskets at the nurse cell-oocyte ring canals [24, 47]. These baskets are only found at the donor side of the ring canals (nurse cells), but never on the recipient side (the oocyte). This localization is established at stages 6-7 and persist to stages 9-10 [24]. These asymmetrically positioned actin filaments can be labeled either by Phalloidin staining (Figure 4A-4B) or by germline-specific expression of LifeAct-TagRFP [48](Figure 4C; Video 8). We quantified the LifeAct-TagRFP signal on both sides of the ring canals connecting the nurse cells and the oocyte, and found a high asymmetry on the nurse cell side over the oocyte side (Figure 4F). This asymmetry of actin fibers at ring canals sharply decreases between nurse cells towards the anterior side of the egg chamber, with the lowest asymmetry at ring canals connecting anterior-most nurse cells (Figure 4E-4F). This level of actin asymmetry correlates well with the directionality of the cargo transport: we observed less directional bias of transport through the ring canals connecting nurse cells where asymmetry of actin fibers is significantly less than the nurse cell-oocyte ring canals (Figure 4G).

**Figure 4.**
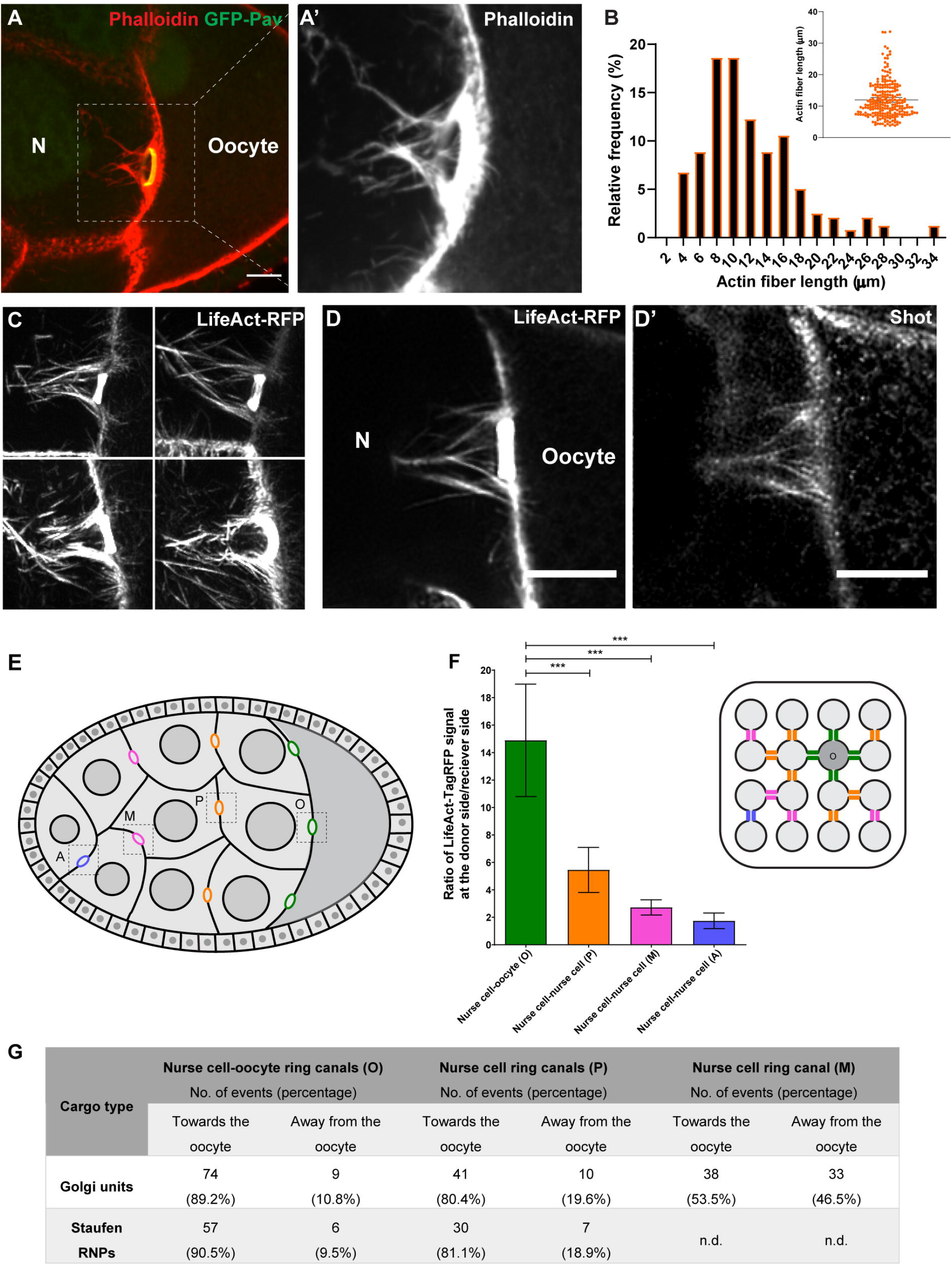
Shot is localized at the asymmetric actin fibers at the ring canals. (A-A’) Rhodamine-conjugated phalloidin staining shows asymmetric actin fibers (the white dashed box) at the ring canal (ring canal inner rim is labeled with GFP-Pavarotti) on the nurse cell side, not at the oocyte side. (B) Quantification of the length of actin fibers on the nurse cell side. The length of four longest actin fibers was measured for each ring canal (59 ring canals from 15 egg chambers). The average actin fiber length on the nurse cell side is 12.0 ± 0.7 μm (mean ± 95% confidence interval). (C) Asymmetric actin fibers, labeled with TagRFP-tagged LifeAct, are seen at all four ring canals connecting nurse cells and the oocyte in the live sample. See also in Video 8. (D-D’) A reprehensive image of Shot antibody staining in a TagRFP-LifeAct-expressing egg chamber. Shot is localized at the asymmetric actin fibers on the nurse cell side of the ring canal, but it is not concentrated in the F-actin core of the ring canal inner rim. (E) A schematic illustration of a *Drosophila* egg chamber at stage 8. Ring canals are categorized depends on its relative distance to the oocyte and are color-coded: (1) nurse cell-oocyte ring canals, directly connected to the oocyte, green, “O”; (2) posterior nurse cell-nurse cell ring canal, having one nurse cell between this ring canal and the oocyte, orange, “P”; (3) middle nurse cell-nurse cell ring canal, having two nurse cells between this ring canal and the oocyte, magenta, “M”; (4) anterior nurse cell-nurse cell ring canal, having three nurse cells between this ring canal and the oocyte, blue, “A”). (F) The asymmetry of actin fibers is quantified as the ratio of LifeAct-TagRFP fluorescence signal at the anterior side to the signal at the posterior side of the ring canals. Mann Whitney tests were performed in following groups: “O” and “P”, p<0.0001 (***); “O” and “M”, p<0.0001 (***);“O” and “A”, p<0.0001 (***). (D) Summary of directionality of two type of cargoes (Golgi units and Staufen RNP particles) at different ring canals. Golgi units are labeled with RFP-Golgi and Staufen RNP particles are labeled with Staufen-SunTag [73]. Number of events are divided into two groups: “towards the oocyte” (moving towards the posterior) and “away from the oocyte” (moving towards the anterior). Highest directionality of both Golgi and Staufen transport was observed at the nurse cell-oocyte ring canals. N, nurse cell; scale bars, 10 μm.

We next examined localization of Shot using immunostaining with a monoclonal antibody against Shot [12]. We found that Shot staining is associated with these asymmetric actin fibers, showing a high level of asymmetry at the nurse cell-oocyte boundary (Figure 4D-D’). This specific localization of Shot at the ring canals implies that Shot controls the transport of cargoes from nurse cells to the oocyte through its interaction with these asymmetric actin filaments.

### Shot organizes microtubules in the ring canals

Having established that Shot is required for directional cargo transport between nurse cells and the oocyte and is asymmetrically localized at the actin fibers on the nurse cell side, we decided to investigate the mechanism by which Shot controls the direction of transport through the nurse cell-oocyte ring canals. As microtubules are found inside the ring canals connecting nurse cells with the oocyte, the transport of organelles and mRNA/proteins to the oocyte have been considered as typical examples of microtubule-dependent motor-driven transport [23-25, 39, 42, 49]. Therefore, we examined whether *shot-RNAi* changes microtubule tracks in the ring canals connecting nurse cells and the oocyte. First, we labeled the microtubules by overexpressing the minus-end binding protein, Patronin [50, 51], and found that, consistent with previous reports [23, 25], microtubules are present at the ring canals between nurse cells and the oocyte, as well as between nurse cells (Figure 5A-A’’). Microtubule distribution is not visibly altered either in the nurse cells or at the ring canals of *shot-RNAi* egg chambers (Figure 5B-B’’). Next we examined the orientation of microtubules running through the ring canals by expressing GFP-tagged EB1 [52]. We found that in control most of the microtubules in the ring canals are oriented with their plus-ends pointing towards nurse cells (Figure 5C-C’’ and 5E; Video 9). In sharp contrast, in *shot-RNAi* mutant there are fewer EB1 comets at the ring canals, and direction of the EB1-GFP comets shows that these microtubules have a mixed orientation (Figure 5D-5E; Video 9).

**Figure 5.**
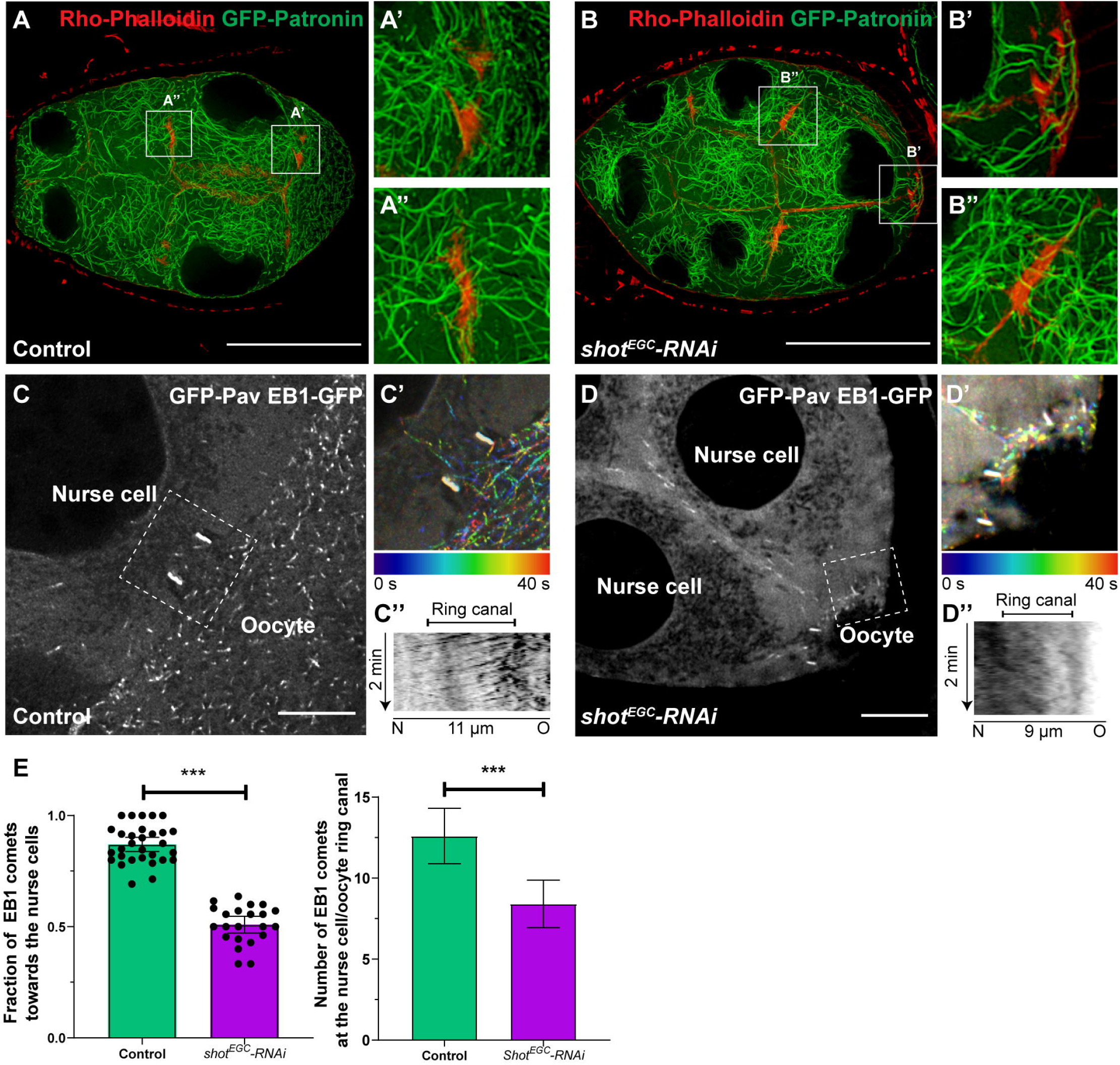
Shot controls microtubule polarity in the ring canal. (A-B) Overall microtubule organization is not affected by *shot* knockdown. (A) In control, microtubules are localized at the ring canals between the nurse cell and the oocyte (A’) and between two nurse cells (A’’). (B) Knockdown of *shot* does not change microtubule distribution at the ring canals between the nurse cells and the oocyte (B’) and between two nurse cells (B’’). Microtubules are labeled by overexpressed GFP-tagged Patronin, and ring canals are labeled with Rho-Phalloidin staining. Scale bars, 50μm. (C-E) Knockdown of *shot* results in a mixed orientation of microtubules in the ring canals. (C) EB1-GFP-labeled microtubule +end comets at the ring canal (labeled by GFP-Pav) connecting the nurse cell and the oocyte in control. (C’) A color-coded hyperstack of the EB1 comet movement of (C). (C’’) Kymograph of EB1 comet movement at the ring canal (the white dashed box in C) in control. (D) EB1-GFP-labeled microtubule +end comets at the ring canal (labeled by GFP-Pav) connecting two nurse cells and the oocyte in *shot-RNAi*. (D’) A color-coded hyperstack of the EB1 comet movement of (D). (D’’) Kymograph of EB1 comet movement at the ring canal (the white dashed box in D) in *shot-RNAi*. (E) Quantification of the fraction of EB1 comets moving through the ring canals towards the nurse cells, and quantification of number of comets at the ring canals in control and *shot-RNAi*. Left, Mann-Whitney test, p-value < 0.0001; right, Mann-Whitney test, p-value < 0.0001. N, nurse cell; O, oocyte; scale bars, 10μm.

Interestingly, by dual labeling with LifeAct-TagRFP and EB1-GFP, we found that microtubule plus-ends tend to grow along the actin fibers of the ring canals on the nurse cell side (Video 10). Therefore, Shot may have a role in favoring microtubule growth along the asymmetric actin fibers at the ring canals, which could facilitate oocyte microtubule plus-ends pass through the ring canal, while preventing nurse cell microtubule plus-ends from entering the ring canal, thus controlling microtubule polarity (Figure 6).

**Figure 6.**
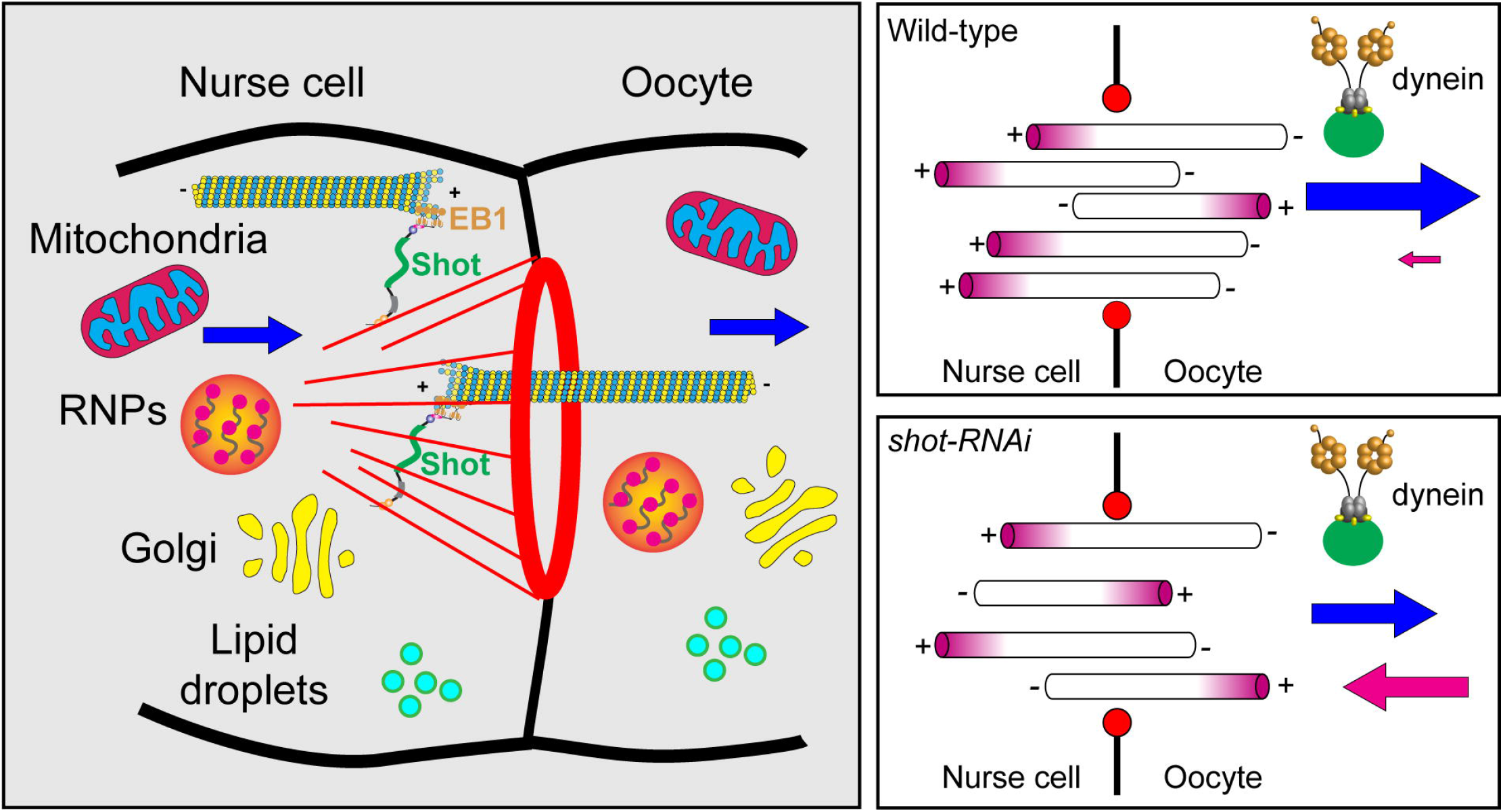
Shot is a gatekeeper at the ring canal for Drosophila oocyte growth. Shot controls microtubule orientation at the ring canal, via regulating the interaction between EB1/microtubule plus-ends and asymmetric actin fibers on the nurse cell side. Therefore, Shot is essential for dynein-dependent transport of various cargoes (including mitochondria, *osk*/Staufen RNPs, Golgi units and lipid droplets) to the oocyte.

The minus-end directed motor cytoplasmic dynein has been proposed to transport organelles and RNP granules to the oocyte [23-25, 32]. To examine whether dynein is required for oocyte growth, we knocked down dynein heavy chain in the ovary by RNAi (*DHC64C-RNAi,* [53]). In order to bypass dynein’s requirement for cell division and oocyte specification, we used the maternal α-tubulin-Gal4 (as previously). We found that that dynein knockdown mimics the “small oocyte” phenotype that we observed in *shot-RNAi* (Supplemental Figure S2G-S2I).

As we observed the difference in the orientation of microtubule tracks at the ring canals between control and *shot-RNAi*, it is highly possible that the microtubule of mixed polarity causes the minus-end directed dynein moving cargoes bidirectionally through the nurse cell-oocyte ring canals, which limits the accumulation of cytoplasmic materials in the oocyte, and eventually stalls the oocyte growth (Figure 6).

Together, we propose that Shot functions as a gatekeeper at the ring canal: Shot favors the uniform microtubule orientation with minus-ends into the oocyte, and allowing cytoplasmic dynein to transport various cargoes from the nurse cells to the oocyte to ensure its rapid growth during oogenesis (Figure 6).

## DISCUSSION

Spectraplakin proteins coordinate and regulate two major cytoskeletal networks, microtubules and actin filaments. *Drosophila* has only a single spectraplakin protein Short stop (Shot) that makes it a perfect model to study the interaction and coordination between microtubules and F-actin. In the biggest cell of the whole animal, the oocyte, microtubules and F-actin are dynamic but precisely arranged throughout the development. In this study, we show that Shot is absolutely required for the oocyte growth. Shot is localized asymmetrically at the actin fibers on the nurse cell side of the ring canals, and controls the microtubule polarity in the ring canals connecting nurse cells and the oocyte. Therefore, Shot directs cytoplasmic transfer of many if not all cargoes produced in nurse cells that are essential for rapid oocyte growth.

### Asymmetric actin baskets at the ring canals

The asymmetric actin baskets at the ring canal start forming in stage 6-7 egg chambers and persist to stage 9-10 before they become indistinguishable from the actin cables formed for nurse cell dumping [24, 47]. From stage 6 to stage 10, the oocyte experiences exponential growth in size [54] caused by unidirectional transport of material from nurse cells to the oocyte. This flow of material precedes massive nurse cell dumping caused by contraction of nurse cells at stages 11-12 [22], and is easy to distinguish from dumping because during the directional transport stage the volume of the oocyte increases but the nurse cells do not shrink in size. Strong correlation between the appearance of these asymmetric actin baskets and the rapid oocyte growth suggests that these actin baskets are involved in directing of transport through the ring canals to the oocyte.

It is not yet clear why the actin baskets are formed asymmetrically at the nurse cell-oocyte border, while the asymmetry is much less prominent in the ring canals between nurse cells. One possible explanation is that actin filaments are organized differently in the nurse cells and in the oocyte. Actin filaments form microvilli originating from the plasma membrane in nurse cells, and are more abundant in a close proximity to the ring canals. Formation of these microvilli depends on the *Drosophila* Profilin homolog, Chickadee, and Fascin homolog Singed [55, 57]. Meanwhile, oocyte has a uniform cortex composed of randomly oriented F-actin and an actin cytoplasmic mesh organized by two actin filament nucleators Cappuccino (*Drosophila* Formin homolog) and Spire (contains of four WASP homology 2 (WH2) domains) [58, 60]. Therefore, albeit in a 16-cell syncytium connected by ring canals, it is likely that different regulation of actin growth leads to asymmetric actin baskets formed only on the nurse cell side, but not on the oocyte side of these ring canals [61].

### Shot guides microtubules at the ring canal

Shot is a large multi-domain cytoskeletal protein that crosslinks two major cytoskeletal components, microtubules and actin filaments, in *Drosophila*. Our results show that Shot is localized at the asymmetric actin fibers at the ring canals between nurse cells and the oocyte. Interestingly, microtubules are required for maintaining these actin baskets at the ring canals. Depolymerization of microtubules in the egg chamber by treatment with microtubule-depolymerizing drug, colchicine, results in disappearance of the actin baskets at the ring canals [24]. It implies that Shot is not just passively localized at the baskets; instead, it plays a more active role in stabilizing and maintaining their structure, probably via its interaction with microtubules and actin filaments.

Our data suggest that Shot plays a role in guiding the microtubule plus-ends along the actin fibers. Microtubules at the ring canals have more plus-ends towards the nurse cells, while knockdown of Shot changes these ring canal microtubules to a mixed orientation. This Shot function is probably dependent on the EB1-interacting SxIP motifs at the C-terminal tail. Studies in *Drosophila* neurons showed that Shot interacts with EB1 protein and F-actin in the growth cone, and thus guilds polymerizing microtubules along actin-structure in the direction of axonal growth [5, 8, 62]. Additionally, Shot has been shown to promote microtubule assembly by recruiting EB1/APC2 at the muscle-tendon junctions [63]. These studies are in an agreement with our model that Shot favors the microtubule polymerization along the asymmetric actin fibers, which in turn allows microtubule plus-ends in the oocyte to enter nurse cells, and meanwhile prevents microtubule plus-ends in the nurse cells from entering the ring canals. Therefore, Shot localization at the asymmetric actin baskets results in the directional bias of microtubule tracks, allowing the minus-end-directed motor dynein to efficiently transfer cytoplasmic contents to the oocyte (Figure 6).

### Intercellular cytoplasmic bridges are conserved across species

In this study, we demonstrate that multiple cargoes, including Golgi units, RNP granules, mitochondria and lipid droplets are transported through the ring canals from the nurse cells to the oocytes, of which the directionality is controlled by the microtubule-actin cross-linker Shot. The ring canal in *Drosophila* egg chambers is not the sole example of cytoplasmic bridges connecting cells and transferring cytoplasm. Multiple organisms ranging from plants to mammals have arrested cytokinesis and maintain the contractile rings as stable cytoplasmic bridges to stay connected between sister cells, both in germline cells and in somatic cells [64]. In *C. elegans* oogenesis, growing oocytes are connected with transcriptionally active germ cells through cytoplasmic bridges, and receive materials from these germ cells, including mitochondria and P-granule components [65]. Remarkably, mouse germ cyst cells also transfer organelles, such as Golgi and mitochondria, to the developing oocyte through ring canals in a microtubule transport-dependent manner [66]. This “sister cell transferring cytoplasm” paradigms in worms and in mice highly resemble the *Drosophila* nurse cell-to-oocyte transport, suggesting it could be an evolutionarily conserved mechanism of cytoplasmic transfer during germline development. This intercellular transfer may present a highly efficient way for the oocyte to acquire essential materials/organelles for its rapid growth.

Altogether, we illustrate that *Drosophila* spectraplakin Shot functions as a gatekeeper at the cytoplasmic canal, and controls the directionality of cytoplasmic transfer from the nurse cells to the oocyte, which ensures the oocyte to have enough cytoplasmic materials during its rapid growth. As spectraplakin family proteins and intercellular cytoplasmic bridges are conserved across species, it is likely that it serves as a universal cytoplasmic transfer mechanism for oocyte growth in higher organisms.

## ACKNOWLEDGEMENTS

We would like to thank Dr. David Glover (Caltech) for *ubi-GFP-Pav* line, Dr. Derek Applewhite (Reed College) for full-length shot cDNA (shot.LA) and *ubi-EB1-GFP* line, Dr. Ferenc Jankovics (Institute of Genetics, Biological Research Centre of the Hungarian Academy of Sciences) for *shot*^Δ*EGC*^ line, Dr. Daniel St Johnston (University of Cambridge) for *mat* α*tub-RFP-*Staufen line, Dr. Vladislav Verkhusha (Albert Einstein College of Medicine) for MoxMaple3 construct, Dr. Michael Welte (University of Rochester) for *UASp-GFP-LD* line, Dr. Uri Abdu (Ben-Gurion University of the Negev) *for UASp-GFP-Patronin* line, Dr. Antoine Guichet (CNRS, Institut Jacques Monod) for *UASp-EB1-GFP* line, the Bloomington *Drosophila* Stock Center (supported by National Institutes of Health grant P40OD018537) for fly stocks, and *Drosophila* Genomics Resource Center (supported by National Institutes of Health Grant 2P40OD01094) for DNA constructs. The Orb 4H8 monoclonal antibody developed by Dr. Paul D. Schedl’s group at Princeton University, and anti-Shot mAbRod1 antibody developed by Dr. Peter A. Kolodziej’s group at Vanderbilt University were obtained from the Developmental Studies Hybridoma Bank, created by the NICHD of the NIH and maintained at The University of Iowa. We also thank all the Gelfand laboratory members for support, discussion, and suggestions. Research reported in this study was supported by the National Institute of General Medical Sciences grants R01GM124029 and R35GM131752 to V.I. Gelfand.

## AUTHOR CONTRIBUTIONS

W.L., M.L., and V.I.G. planned and designed the research. W.L. and M.L. conducted experiments and data analysis; W.L. and V.I.G. wrote the manuscript.

## Materials and methods

### Plasmid constructs

The oligos of shot^ABD^-shRNA (agtTGCGCGATGGTCACAATTTACtagttatattcaagcataGTAAATTGTGACCATCGCGCA gc) and shot^EGC^-shRNA (agtCCGGAAAATGGATAAGGATAAtagttatattcaagcataTTATCCTTATCCATTTTCCGGg c) were synthesized and inserted into the pWalium22 vector (*Drosophila* Genomics Resource Center, Stock Number #1473, 10XUAS)[67] by NheI(5’)/EcoRI(3’). *shot*^*ABD*^-*RNAi* targeting sequences: TGCGCGATGGTCACAATTTAC; *shot^EGC^*-RNAi targeting sequences: CCGGAAAATGGATAAGGATAA.

MoxMaple3 was amplified by PCR from the pmCherry-T2A-moxMaple3 (Addgene Plasmid #120875) [43] and inserted into the pUASp by SpeI (3’)/EcoRI (3’); mitochondria targeting probe, human Cox8a (mitochondrial cytochrome c oxidase subunit 8A) (1-29 residues, MSVLTPLLLRGLTGSARRLPVPRAKIHSL) (atgtccgtcctgacgccgctgctgctgcggggcttgacaggctcggcccggcggctcccagtgccgcgcgccaagatccattcgttg) was synthesized and inserted into pUASp-MoxMaple3 by KpnI(5’)/SpeI(3’).

All the constructs were sent to BestGene for injection: PhiC31-mediated integration (*UAS-shot^ABD^-RNAi* and *UAS-shot*^*EGC*^-*RNAi in pWalium22 vectors*, at attP2 site) and P-element insertion (*pUASp-Mito-MoxMaple3*).

### *Drosophila* genetics

Fly stocks and crosses were maintained on standard cornmeal food (Nutri-Fly® Bloomington Formulation, Genesee, Cat #: 66-121) supplemented with dry active yeast at room temperature (~24– 25°C), The following fly stocks were used in this study: *hs-FLP^[12]^* (X, Bloomington *Drosophila* Stock Center #1929 [68]); *FRTG13* (II, Bloomington *Drosophila* stock center # 1956); *FRTG13 shot^[3]^* (Bloomington *Drosophila* Stock Center # 5141 [69]); *FRTG13 ubi-GFP.nls* (II, Bloomington *Drosophila* Stock Center # 5826); *shot*^Δ*EGC*^ (from Dr. Ferenc Jankovics, Institute of Genetics, Biological Research Centre of the Hungarian Academy of Sciences [19]); *mat* α*tub-Gal4^[V37]^* (III, Bloomington *Drosophila* Stock Center #7063); *ubi-GFP-Pav* (from Dr. David Glover, Caltech [35]); *shot^Rod^-RNAi* (TRiP.GL01286, attP2, III, Bloomington *Drosophila* Stock Center # 41858), *UASt-shot.L(A)*Δ*rod-GFP* (Bloomington *Drosophila* Stock Center # 29040 [7]); *shot*^Δ*ABD*^ (aka *shot^[k03010]^*, *shot^[kakP1]^*; Bloomington *Drosophila* Stock Center #10522 [6, 7, 26, 70]); *nos-Gal4-VP16* (III [71, 72]); *UASp-LifeAct-TagRFP* (II, 22A, Bloomington *Drosophila* Stock Center # 58713); *UASp-LifeAct-TagRFP* (III, 68E, Bloomington *Drosophila* Stock Center # 58714); *UASp-RFP-Golgi* (II, Bloomington *Drosophila* Stock Center # 30908, aka *UASp-GalT-RFP* [38]); *mat* α*tub-RFP-Staufen* (X, from Dr. Daniel St Johnson [40]); *UASp-GFP-LD* (II, from Dr. Michael Welte [46]); *UASp-GFP-Patronin* (II) (from Dr. Uri Abdu, Ben-Gurion University of the Negev [30, 51, 73]); *UASp-EB1-GFP* (II, from Dr. Antoine Guichet [52]); *ubi-EB1-GFP* [53, 74]; *UASp-F-Tractin-tdTomato* (II, Bloomington stock center #58989, [75]); *UAS-Dhc64C-RNAi* (TRiP.GL00543, attP40, II, Bloomington *Drosophila* Stock Center # 36583)[53]; *UASp-Staufen-SunTag* (III, [73]). The following fly stocks were generated in this study: *UAS-shot*^*ABD*^-*RNAi* (in pWalium22 vector, inserted at attP2, III)*; UAS-shot^EGC^-RNAi* (in pWalium22 vector, inserted at attP2, III)*; UASp-Mtio-MoxMaple3* (II).

### Induction of germline clones of *shot*^*[3]*^ and *shot*^Δ*EGC*^

A standard recombination protocol was performed between *FRTG13* and *shot*^Δ*EGC*^. FRTG13 *shot^[3]^*/*CyO* or *FRTG13 shot*^Δ*EGC*^*/CyO* virgin female flies were crossed with males carrying *hs-flp*^*[12]*^*/y; FRTG13 ubi-GFP.nls/CyO*. From these crosses, young pupae at day 7 and day 8 AEL (after egg laying) were subjected to heat shock at 37 °C for 2 hours each day. Non CyO F1 females were collected 3-4 day after heat shock and fattened with dry active yeast overnight before dissection for Orb staining.

### Immunostaining of *Drosophila* oocytes

A standard fixation and staining protocol was used [72, 76]. Samples were incubated with mouse monoclonal anti-Orb antibody (Orb 4H8, Developmental Studies Hybridoma Bank, 1:5) or mouse monoclonal anti-Shot antibody (shot mAbRod1, Developmental Studies Hybridoma Bank, 1:5) at 4°C overnight, washed with 1XPBTB (1XPBS + 0.1% Triton X-100 + 0.2% BSA) five times for 10 min each time, incubated with FITC-conjugated or TRITC-conjugated anti-mouse secondary antibody (Jackson ImmunoResearch Laboratories, Inc; Cat# 115-095-062 and Cat# 115-025-003) at 1:100 at room temperature (24~25°C) for 4 h, and washed with 1XPBTB five times for 10 min each time. Some samples were stained with Rhodamine-conjugated phalloidin (1:5000) for 30 min before mounting. Samples were imaged either on a Nikon A1plus scanning confocal microscopy with a GaAsP detector, and a 20× 0.75 N.A. lens using Galvano scanning, or on a Nikon Eclipse U2000 inverted stand with a Yokogawa CSU10 spinning disk confocal head and a 40× 1.30 NA oil lens using an Evolve EMCCD, both controlled by Nikon Elements software. Images were acquired every 1 μm/step for whole ovariole imaging or 0.5 μm/step for individual egg chambers in z stacks.

### Live imaging of *Drosophila* egg chamber

Young mated female adults were fed with dry active yeast for 16~18 hours and then dissected in Halocarbon oil 700 (Sigma-Aldrich) as previously described [30, 73, 76]. Fluorescent samples were imaged using Nikon W1 spinning disk confocal microscope (Yokogawa CSU with pinhole size 50um) with Photometrics Prime 95B sCMOS Camera, and a 40 x 1.30 N.A. oil lens or a 40X 1.25 N.A. silicone oil lens, controlled by Nikon Elements software.

### Labeling of microtubules by GFP-Patronin in *Drosophila* egg chambers

Ovaries from flies expressing GFP-Patronin under *maternal* α*tub-Gal4*^*[V37]*^ (with or without the *UAS-shot*^*EGC*^-RNAi) were dissected and fixed in 1XPBS +0.1%Triton X-100 +4% EM-grade formaldehyde for 20 min on the rotator; briefly washed with 1XPBTB five times and stained with Rhodamine-conjugated phalloidin for 30 min before mounting. Samples were imaged using Nikon W1 spinning disk confocal microscope (Yokogawa CSU with pinhole size 50um) with Photometrics Prime 95B sCMOS Camera, and a 40 × 1.30 N.A. oil lens, controlled by Nikon Elements software. Images were acquired every 0.3 mm/step in z stacks and 3D deconvoluted using Richardson-Lucy iterative algorithm provided by Nikon Elements. A maximum intensity projection of 0.6 μm z-stack sample (3 slices in the z-stacks) was used to present the microtubule distribution in each genotype.

### Quantification of cargo transport direction and microtubule polarity

Kymographs were created along a ~3.7 μm-width line (for cargo transport) or a 3.5 μm ~5.0 μm - width line (for microtubule polarity) from the nurse cell to the oocyte through the ring canals (labeled by GFP-Pav or F-Tractin-tdTomato). Cargo movement direction and microtubule polarity were manually quantified based on these kymographs.

