## Supplementary Figures for "Short stop is a gatekeeper at the ring canals of *Drosophila* ovary"

**Supplementary figures and legends**

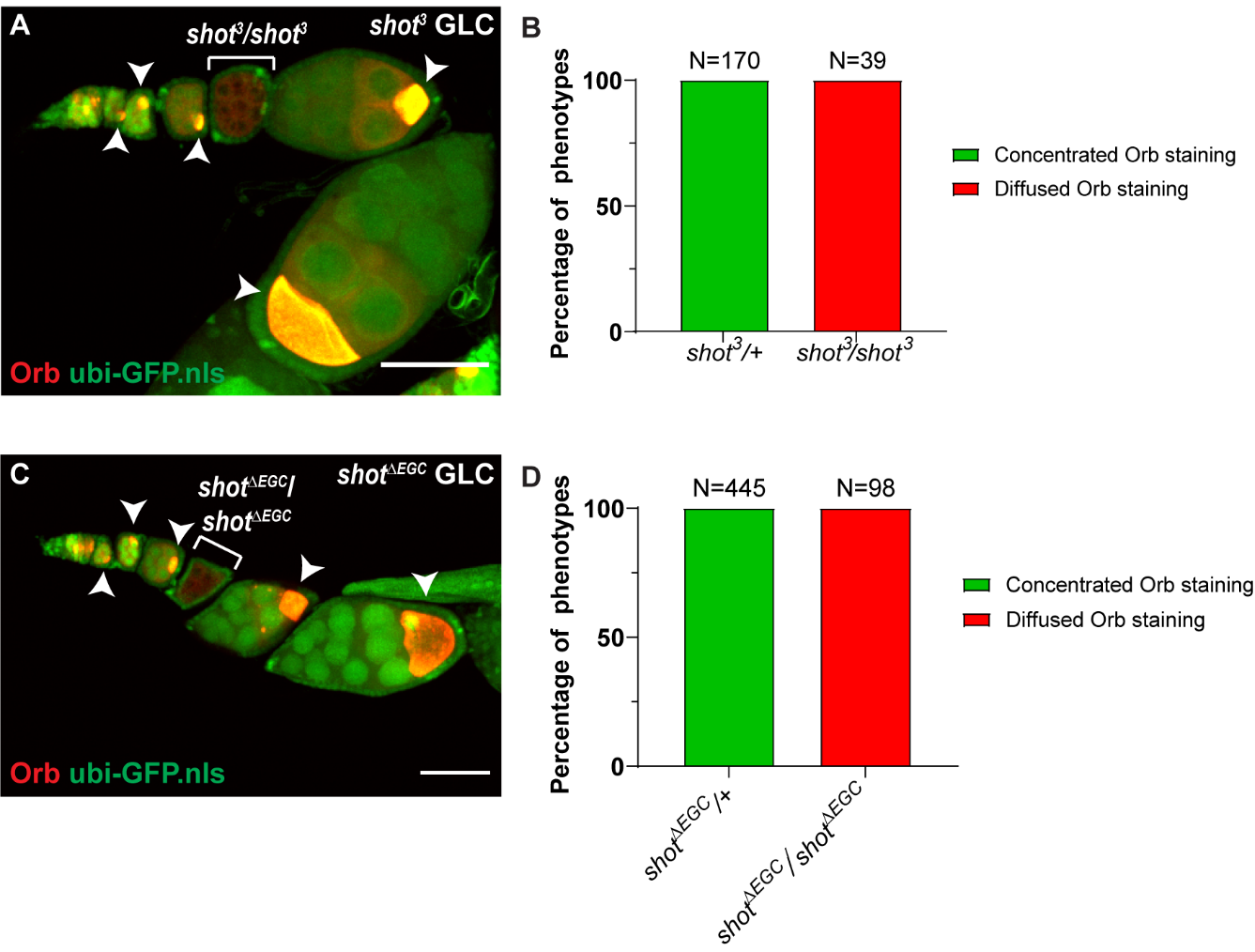

**Supplementary Figure 1. Shot is required for oocyte specification.**

(A) A germline clone that is homozygous of *shot^[3]^*, a strongest loss-of-function allele, fails to specify the oocyte in the egg chamber, shown by the lack of concentrated Orb staining (the white bracket). *shot^[3]^* heterozygous egg chambers have normal oocyte specification (white arrowheads). (B) Summary of the Orb staining phenotypes in *shot^[3]^* heterozygous and homozygous egg chambers.

(C) A germline clone that is homozygous of *shot^∆EGC^* fails to specify the oocyte in the egg chamber, shown by the lack of concentrated Orb staining (the white bracket). *shot^∆EGC^* heterozygous egg chambers have normal oocyte specification (white arrowheads). (B) Summary of the Orb staining phenotypes in *shot^∆EGC^* heterozygous and homozygous egg chambers.

Scale bars, 50 µm.

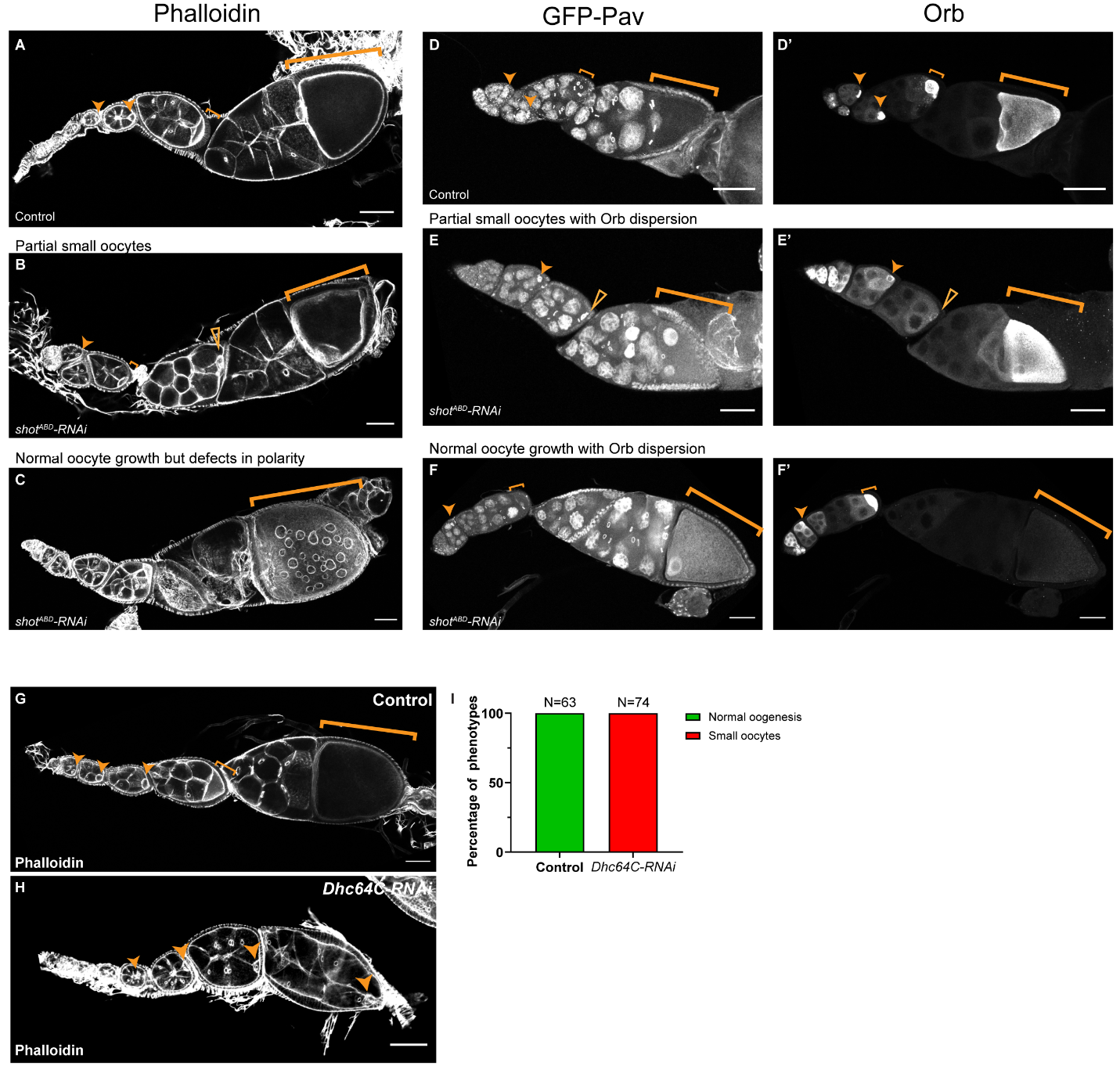

**Supplementary Figure 2. Oogenesis defects in *shot-RNAi* and in *dynein-RNAi*.**

(A-C) In addition to complete small oocyte phenotype (shown in Figure 1 and Video 1), *shot-RNAi* also display partial small oocytes (B, pointed by the open arrowhead), and normal oocyte growth with polarity defects (C, highlighted by the bracket), compared to normal oogenesis in control (A). Percentages of these phenotypes are summarized in Figure 1F.

(D-F’) *shot-RNAi* also display partial small oocytes with Orb dispersion (E-E’, pointed by the open arrowhead), and normal oocyte growth with Orb dispersion (F-F’, highlighted by the big bracket), compared to normal oogenesis with Orb concentration in control (D-D’). Percentages of these phenotypes are summarized in Figure 1G.

(G-H) Representative images of Rhodamine-conjugated phalloidin staining in control (*mat αtub-Gal4^[V37]^/+*) and *dynein-RNAi* (*UAS-Dhc64C-RNAi*/+; *mat αtub-Gal4^[V37]^/+*). Normal oocyte growth in control, shown by orange arrowheads and brackets (G); oocytes remained small in *dynein-RNAi*, shown by orange arrowheads (H).

(I) Summary of oocyte phenotypes in control and *Dhc64C-RNAi*.

Scale bars, 50 µm.

**Video legends**

**Video 1. Shot knockdown leads to oocyte growth defects.** Representative Z-stack images of phalloidin staining in control (*yw;* *mat αtub-Gal4^[V37]^*/+), *shot^ABD^-RNAi* (*yw;* *mat αtub-Gal4^[V37]^*/*UAS-shot^ABD^-RNAi*), and *shot^EGC^-RNAi* (*yw;* *mat αtub-Gal4^[V37]^*/*UAS-shot^EGC^-RNAi*). Scale bars, 50 µm.

**Video 2. Shot knockdown does not affect initial oocyte specification but causes to loss of oocyte identity over-time.** Representative Z-stack images of Orb staining with GFP-Pav in control (*yw*; *ubi-GFP-Pav*/+; *mat αtub-Gal4^[V37]^*/+), *shot^ABD^-RNAi* (*yw*; *ubi-GFP-Pav*/+; *mat αtub-Gal4^[V37]^*/ *UAS-shot^ABD^-RNAi*), and *shot^EGC^-RNAi* (*yw*; *ubi-GFP-Pav*/+; *mat αtub-Gal4^[V37]^*/*UAS-shot^EGC^-RNAi*). Scale bars, 50 µm.

**Video 3. The directionality of Golgi transport is disrupted in *shot-RNAi* mutant.** Golgi units are labeled with RFP-Golgi (GalT-RFP) and ring canals are labeled with GFP-tagged kinesin-6/Pavarotti in control (*yw; UASp-RFP-Golgi/ubi-GFP-Pav; mat αtub-Gal4^[V37]^/+*) and in *shot-RNAi* (*yw; UASp-RFP-Golgi/ubi-GFP-Pav; mat αtub-Gal4^[V37]^/UAS-shot^EGC^-RNAi*). Egg chamber is oriented as the anterior to the left and the posterior to the right. Time lapses images were acquired every 5 sec for 5 min; scale bars, 50 µm.

**Video 4. Staufen particles are transported more often from the oocyte to the nurse cells in *shot-RNAi* mutant.** Staufen particles are labeled with RFP-Staufen and ring canals are labeled with GFP-tagged kinesin-6/Pavarotti in control (*w, mat αtub-RFP-Staufen/yw; ubi-GFP-Pav/+; mat αtub-Gal4^[V37]^/+*) and in *shot-RNAi* (*w, mat αtub-RFP-Staufen/yw; ubi-GFP-Pav/+; mat αtub-Gal4^[V37]^/UAS-shot^EGC^-RNAi* ). Egg chamber is oriented as the anterior to the left and the posterior to the right. Time lapses images were acquired every 5 sec for 5 min; scale bars, 50 µm.

**Video 5. Photoconverted mitochondria move towards the oocyte in control.** Mitochondria are labeled with Mito-MoxMaple3 and a subset of mitochondria were photoconverted from green to red by local UV exposure, either in the posterior most nurse cells or in the oocyte. The ring canals are labeled with GFP-tagged kinesin-6/Pavarotti (*yw; ubi-GFP-Pav/UASp-Mito-MoxMaple3; mat αtub-Gal4^[V37]^/+*) and shown as a pair of orange arrowheads in the zoom-in areas. Egg chamber is oriented as the anterior to the left and the posterior to the right. A max projection of a total 4 µm Z stack was shown for both green and red channel. Time lapses images were acquired every 30 sec for 20 min; scale bars, 50 µm.

**Video 6. *shot-RNAi* changes the direction of mitochondria movement between the nurse cells and the oocyte.** Mitochondria are labeled with by Mito-MoxMaple3 after global photoconversion and ring canals are labeled with GFP-tagged kinesin-6/Pavarotti in control (*yw; ubi-GFP-Pav/UASp-Mito-MoxMaple3; mat αtub-Gal4^[V37]^/+*) and in *shot-RNAi* (*yw; ubi-GFP-Pav/UASp-Mito-MoxMaple3; mat αtub-Gal4^[V37]^/UAS-shot^EGC^-RNAi*). Egg chamber is oriented as the anterior to the left and the posterior to the right. Time lapses images were acquired every 5 sec for 5 min; scale bars, 50 µm.

**Video 7. Lipid droplet transport becomes bidirectional between nurse cells and the oocyte in *shot-RNAi* mutant.** Lipid droplets are labeled with GFP-tagged lipid droplet domain of *Drosophila* protein Klar (GFP-LD) and ring canals are labeled with tdTomato-tagged actin-binding domain of rat inositol triphosphate 3-kinase (F-tractin-tdTomato) in control (*yw; UASp-F-tractin-tdTomato/UASp-GFP-LD; mat αtub-Gal4^[V37]^/+*) and in *shot-RNAi* (*yw; UASp-F-tractin-tdTomato/UASp-GFP-LD; mat αtub-Gal4^[V37]^*/ *UAS-shot^EGC^-RNAi*). Egg chamber is oriented as the anterior to the left and the posterior to the right. Time lapses images were acquired every 5 sec for 5 min; scale bars, 50 µm.

**Video 8. Asymmetric actin fibers at the ring canals between nurse cells and the oocyte.** F-actin is labeled with LifeAct-TagRFP in the live sample (*w*; *UASp-LifeAct-TagRFP*/+; *nos-Gal4-VP16*/+) and asymmetric actin fibers are visible at all four ring canals connecting four nurse cells with the same oocyte (indicated by the shaded boxes). Scale bar, 10 µm.

**Video 9. Shot controls microtubule orientation at the nurse cell-oocyte ring canals.** Microtubule plus-ends are labeled with EB1-GFP and ring canals are labeled with GFP-tagged kinesin-6/Pavarotti in control (*yw*; *UASp-EB1-GFP*/*ubi-GFP-Pav*; *mat atub-Gal4^[V37]^*/+) and in *shot-RNAi* (*yw*; *UASp-EB1-GFP*/*ubi-GFP-Pav*; *mat atub-Gal4^[V37]^*/*UAS-shot^EGC^-RNAi*). N, nurse cell; O, oocyte; time lapses images were acquired every 2 sec for 2 min; scale bars, 10 µm.

**Video 10. Microtubule plus-ends grow along the asymmetric actin fibers on the nurse cell side in control.** Microtubule plus-ends are labeled with EB1-GFP and F-actin is labeled with LifeAct-TagRFP in control (*w*; *UASp-LifeAct-TagRFP*/+; *nos-Gal4-VP16*/*ubi-EB1-GFP*). Either focused on the nurse cell-oocyte ring canal cross-section (first half) or focused on asymmetric actin fibers slightly above the ring canal cross-section (second half). Time lapses images were acquired every 2 sec for 2 min; scale bars, 10 µm.
